# Combining cancer chemotherapeutics with bacterial DNA repair inhibitors to develop novel antimicrobials

**DOI:** 10.1101/2023.03.17.532951

**Authors:** Lorenzo Bernacchia, Arya Gupta, Antoine Paris, Alexandra A. Moores, Neil M Kad

**Author notes:** These authors contributed equally to this work (Lorenzo Bernacchia, Arya Gupta). Creating adjuvant antimicrobials.

## Abstract

Cancer chemotherapeutics kill rapidly dividing cells, which includes cells of the immune system. The resulting neutropenia predisposes patients to infection, which delays treatment and is a major cause of morbidity and mortality. Here we have exploited the cytotoxicity of the anti-cancer compound cisplatin to screen for FDA-approved drugs that impair bacterial nucleotide excision DNA repair (NER), the primary mechanism bacteria use to repair cisplatin lesions. Five compounds have emerged of which three possess ideal antimicrobial properties including cell penetrance, specific activity for NER, and the ability to kill a multi-drug resistant clinically relevant *E. coli* strain. Targeting NER offers a new therapeutic approach for infections in cancer patients by combining antimicrobial activity with cancer chemotherapy.

## Introduction

The fundamental therapeutic approach for cancer chemotherapy is to target fast replicating cells by virtue of their need to replicate DNA (1). However, this approach causes neutropenia by off-target killing of circulating immune cells (2). Coupled with the chemotherapy-induced degradation of physical barriers such as mucous membranes, pathogen penetration is also enhanced (3, 4), further contributing to bacterial infection, the second most common cause of death in cancer patients (5). Therefore, the deployment of antimicrobials is required, these target a series of cellular processes ranging from cell wall synthesis to protein synthesis and DNA metabolism (6). However, prolonged exposure and drug overuse has led to the development of antimicrobial resistance through de novo mutation or gene swaps (7). Despite expanding development and approval of pharmaceuticals (8), antimicrobial development has lagged behind other treatments (9). This slow research pipeline means current antimicrobials are losing effectiveness against new microbial variants, exacerbating the need to develop new antimicrobial drugs (10).

The first platinum based anti-cancer drug, cisplatin (cis-Diaminodichloroplatinum, CIS), was discovered fortuitously to inhibit cell division in *Escherichia coli* (11), stalling replication through the formation of DNA adducts with inter- and intra-strand crosslinks (12). Since its target is DNA, it was subsequently found to similarly inhibit human cell proliferation and therefore was exploited to treat a variety of cancer types (13). In both bacterial and mammalian cells, cisplatin adducts are repaired by the specific activity of enzymes in the nucleotide excision DNA repair (NER) pathway (14, 15). Here the similarities end, bacterial NER uses fewer enzymes and has little homology to its human counterparts (16). In bacteria, NER removes a variety of damage types including cisplatin adducts (17) but is primarily deployed to resolve UV-induced DNA damage. It begins with recognition and verification of DNA distorting lesions by UvrA_2_UvrB_2_, followed by recruitment of an endonuclease (UvrC) that nicks the DNA on the same strand either side of the lesion. This damage-containing oligonucleotide is removed by a helicase (UvrD), before DNA pol I restores the correct DNA (18). Therefore, with impaired NER, bacteria would not be able to repair the damage caused by cisplatin during cancer chemotherapy (14, 19).

Inhibition of NER alone does not kill bacteria (19, 20), therefore, in this study, we identified a series of NER inhibitors by screening a library of FDA-approved compounds that kill *E. coli* only in the presence of a sub-lethal dose of cisplatin; since cisplatin causes DNA damage that is normally repaired by NER. We have further triaged the pool of hits using a series of *in vivo, in vitro, in silico* and single-molecule assays, to confirm the mechanism of action for our lead candidates as inhibition of NER. These findings represent a new mode for antimicrobial action as an adjuvant to cisplatin. We anticipate that these compounds could be administered directly to patients receiving cisplatin-based cancer chemotherapy, thereby protecting them from chemotherapy-induced bacterial infection. To provide initial confirmation that these drug combinations may be useful in patients, we have successfully verified the activity of a subset of these compounds against a multidrug resistant clinical isolate of *E. coli*, responsible for the majority of the hospital acquired infections.

The drugs repurposed in this study offer a significant step forward in the battle against co-infection during cancer treatment, which leads to delays in chemotherapy treatment, and directly risks patients’ health. In addition, by defining bacterial NER as a new drug target, this opens the door to adjuvant antimicrobials that work alongside DNA damaging agents, for wider application against multi-drug resistant bacteria.

## Results

### Screening for growth inhibitors in *E. coli*

The screening protocol used identifies FDA-approved compounds that inhibit the viability of *E. coli* in the presence of cisplatin. These compounds undergo a series of further tests to narrow down the mechanism of action as NER inhibition (Figure 1A). To ensure that drug efflux is not a barrier for drug action, thereby maximising the number of hits from our screen of compounds, we created a drug efflux pump (*tolC*) knockout strain of *E. coli* MG1655 (MG1655 Δ*tolC*). However, to understand the role of efflux, all screens were performed in parallel with WT MG1655 and MG1655 Δ*tolC*. The concentration of cisplatin (4 μg/mL) used in the screen was just below the minimal inhibitory concentration (MIC) we recently determined for these strains (19), and all FDA-approved compounds were used at 20 μM. Growth inhibition was determined through the colorimetric resazurin assay (Figure 1B), which relies on active metabolism to convert the blue coloured resazurin to pink resorufin (20). A screen of 2731 compounds revealed 172 potential NER inhibitors. To provide further certainty that we were targeting NER, we subsequently screened these hits using UV exposure. NER is the primary mechanism bacteria use to protect from UV-irradiation, therefore, we exposed both our *E. coli* strains to a sub-MIC dosage of 75 J/m^2^ at 254 nm (20) before introducing the reduced panel of compounds in the absence of cisplatin. This second step resulted in 34 hits (in MG1655 and MG1655 Δ*tolC*, combined). Based on availability we proceeded to further characterise the best 5 of these hits. Among these, two were directly evaluated for their ability to inhibit UvrA binding to DNA at the single molecule level. The final three lead compounds possessed the ability to kill bacteria with cisplatin and evade the drug efflux pump TolC.

**Figure 1:**
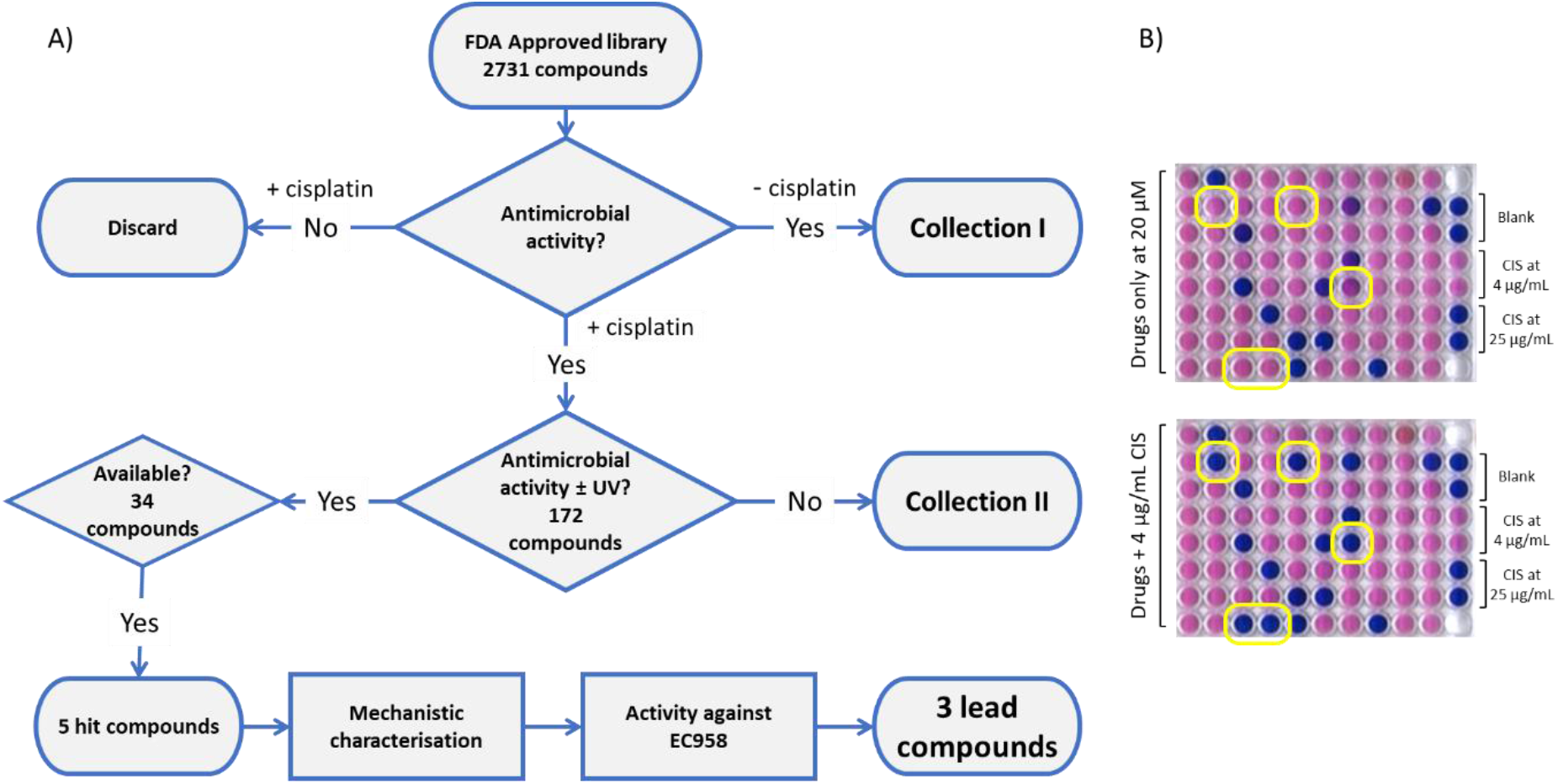
Screening pipeline. Phenotypic screening of FDA approved compounds was performed in the presence of cisplatin using *E. coli* strains MG1655 and the efflux pump knock-out MG1655 Δ*tolC*. The latter was used to increase the search area for active compounds. **A)** Shows a schematic of the screening strategy, starting at finding compounds with antimicrobial activity in the presence of cisplatin and then confirming their activity towards NER using a series of mechanistic assays. Activity against the clinical isolate EC958 identified 3 lead compounds from the original 2731. Collections I and II include a number of potential antimicrobials for future exploration. **B)** Growth inhibition assays in the absence (top) and presence (bottom) of cisplatin. These assays use the colour change of resazurin to indicate bacterial growth (pink) or its inhibition (blue). The appearance of new blue wells in the bottom plate indicates drug activity only in the presence of cisplatin. Dual replicate controls are shown on the right lane (top to bottom) for no drug, sub-lethal cisplatin dose with no drug but containing 2.5% DMSO, and a lethal dose of cisplatin.

### Inhibitor synergy with cisplatin

We firstly established the toxicity of the compounds without cisplatin for both WT MG1655 and MG1655 Δ*tolC*, using resazurin survival assays. With this information, we were able to define the range over which to perform 2-dimenisonal survival assays also known as checkerboards.

An example checkerboard assay is shown in Figure 2A, here the bottom row corresponds to the MIC for cisplatin (the last spot that is blue i.e., 12.5 mg/L) and the leftmost column corresponds to the MIC for the drug (6.25 mg/L). As the drug concentration is raised (right to left in columns) the MIC for cisplatin markedly drops to a maximum effect close to the drug MIC (yellow arrow), this indicates the drug and cisplatin cooperatively inhibit bacterial growth. Each step is a two-fold change in concentration; therefore, the yellow arrow indicates a 16-fold reduction in cisplatin MIC. Similarly, increasing cisplatin (from bottom to top) identifies the maximum cooperative effect on drug dosing (green arrow), for 9-aminoacridine this corresponds to an 8-fold reduction in MIC. The maximum reduction in MIC for cisplatin or drug is shown in Figures 2B & C. All the compounds tested in MG1655 Δ*tolC* (Figure 2B) showed a minimum two-fold decrease in MIC due to drug/cisplatin cooperativity. Figure 2C shows the effects of the compounds that showed activity in TolC-containing WT MG1655; Mitoxantrone, 9-aminoacridine and Pirarubicin all showed 2-fold or greater increases in MIC. As with Δ*tolC* 9-aminoacridine once again showed the strongest effects with an increase in antibacterial activity of 8-fold and increase in cisplatin activity of 4-fold in WT MG1655.

**Figure 2:**
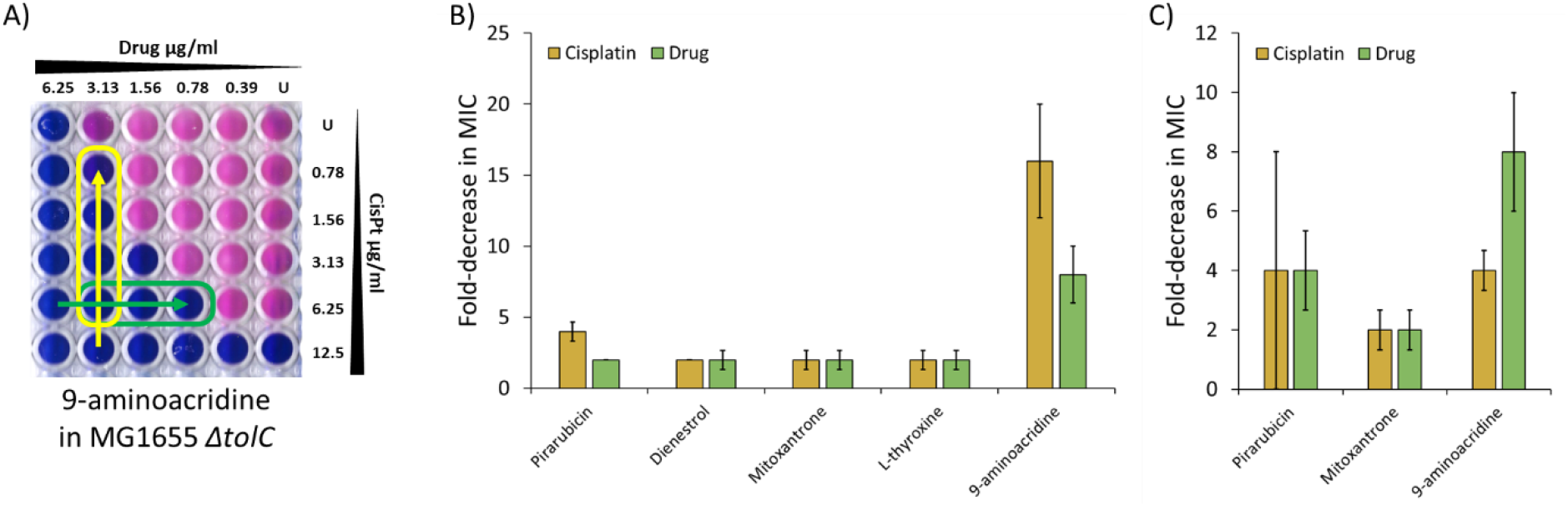
Inhibitory activity of selected hits against MG1655 Δ*tolC* and MG1655 in the presence and absence of cisplatin. **A)** A representative checkerboard assay plate for 9-aminoacridine using MG1655 Δ*tolC* in combination with cisplatin, drug concentration is decreased left to right and [cisplatin] decreases bottom to top. The yellow arrow indicates the greatest decrease in MIC for cisplatin (16-fold) and for the drug (8-fold) this is shown as the green arrow (U = untreated sample). **B)** Bar chart representation of the median fold decrease in MIC for MG1655 Δ*tolC* when the drug and cisplatin were combined (as shown by the arrows in figure A). **C)** Same as (B) but for MG1655. Data points are derived from three independent replicates and error bars are the standard error of the mean.

### Validating NER as the inhibitor target

The above data clearly show that the final set of inhibitors work in combination with cisplatin to inhibit bacterial growth. However, to confirm the mechanism of action we performed a series of studies directly testing efficacy against NER *in vitro* and *in vivo*.

Firstly, we tested for DNA incision in the presence of drug *in vitro*. This crucial step occurs after damage recognition by UvrAB and precedes the resolution aspects of repair and is therefore highly specific for NER. The standard approach for testing incision uses gel-based incision assays (21), however, these are not scalable to high throughput screening and are poorly quantitative. Therefore, we developed a new fluorescence based assay for incision (Figure 3A), which is less prone to photobleaching than another recently developed method (22). A complementary oligonucleotide pair with a 3’ Cy5 on one strand and 5’ black-hole quencher (BHQ) on the other is minimally fluorescent. By placing a fluorescein adducted thymidine 14 nt away, but on the same strand as the Cy5, results in an NER-based incision 10 nt from the Cy5-strand end. This leads to the 10 nt fragment leaving the duplex and an increase in fluorescence (Figure 3A). We expressed and purified UvrA, UvrB and UvrC to quantify the incision reaction using this fluorescence-based assay (Figure 3B), but in parallel confirmed the validity of this approach using a standard gel-based assay (Figure 3C). Pirarubicin (max inhibition of ∼97% after 0.5h of incubation), Mitoxantrone (max inhibition of ∼82% after 16h of incubation) and 9-aminoacridine (max inhibition of ∼49% after 2.5h of incubation) exhibited significant reduction in NER activity, whereas Dienestrol (max inhibition of ∼25% after 16h of incubation) showed only partial inhibition of the pathway. L-thyroxine did not show inhibition in the fluorescence assay, but in the gel-based assay, when incubated for a shorter period (15 minutes), did show inhibition (Figure 3C marked †).

**Figure 3:**
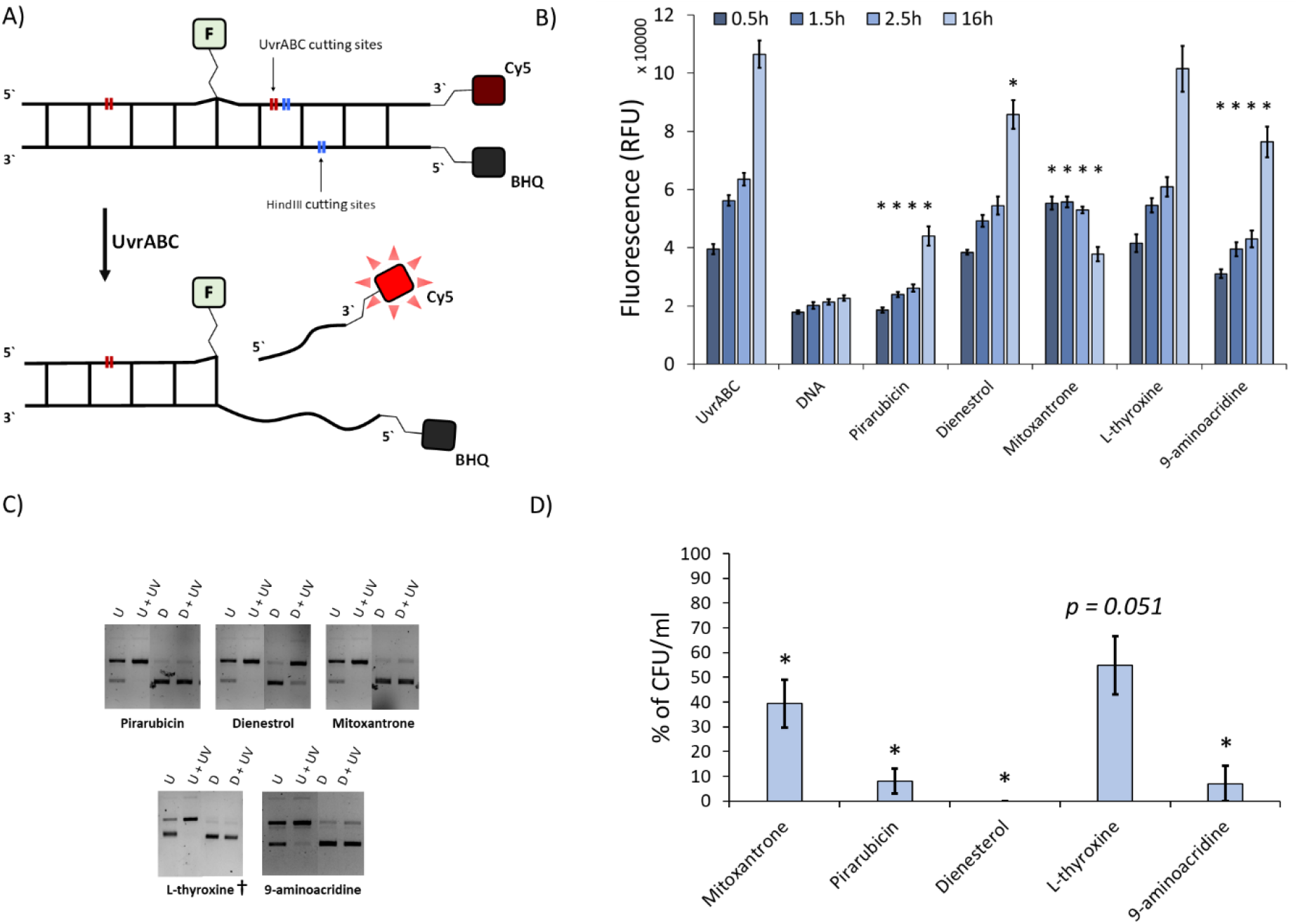
*In vitro* and *in vivo* tests for nucleotide excision repair inhibition. **A)** Schematic representation of the fluorescence incision assay to assess inhibition of NER activity using one oligonucleotide with an engineered damaged sites (F = fluorescein) and a reporter (Cy5 = fluorophore); the second complementary oligonucleotide possessed a black hole quencher (BHQ) to quench the Cy5 fluorescence until the top oligonucleotide is nicked by the NER system proteins UvrA, UvrB and UvrC (UvrABC). **B)** Results from the fluorescence incision assay (A). UvrABC is the control with no drug, and DNA has no drug or UvrABC. The progress was checked at the time points indicated and error bars represent the standard error of the mean. * = p ≤ 0.05 (n ≥ 4 replicates) compared with UvrABC. **(C)** Confirmation of the fluorescence assay with a classical gel-based incision assay demonstrating the inhibition of NER, U is undamaged pUC18, D is the assay in the presence of drug and UV indicates the plasmid is damaged with 200 J/m^2^ UVC (data derives from ≥2 independent replicates) † = 15 minutes incubation. **(D**) Inhibition of plasmid DNA repair *in vivo*. The percent recovery of transformants of pUC18 DNA carrying ampicillin resistance when damaged with 200 J/m^2^ UVC (n = 3) after plating onto ampicillin agar is shown on a scale relative to repair-efficient controls (See figure S1 and S2). Although L-thyroxine substantially reduced repair activity, it remained on the borderline of statistical significance (p = 0.051). Error bars represent the standard error of the mean.

To provide a second test that the drugs were targeting NER, we transformed bacteria with a UVC damaged pUC18 plasmid (254 nm at 200 J/m^2^), carrying the ampicillin selection marker. We reasoned that inhibition of the NER pathway would prevent recovery of transformants on selective agar. As expected, all of the compounds impaired recovery (Figure 3D).

### Drug interactions with the molecules of NER

Having validated that the 5 compounds inhibit NER activity, we sought to investigate how these function on a molecular scale. Upon locating damage UvrA hydrolyses ATP, leading to the loading of UvrB (23–26). We directly measured the rate of purified UvrA’s ATP turnover with and without DNA using an *in vitro* NADH-linked assay (20). When UvrA was incubated with 20 µM of each shortlisted compound, four were found to significantly affect the ATPase (Figure 4A). Among those, Pirarubicin, Mitoxantrone, Dienestrol and L-thyroxine all inhibited the ATPase, with the latter two drugs having the strongest effect.

**Figure 4:**
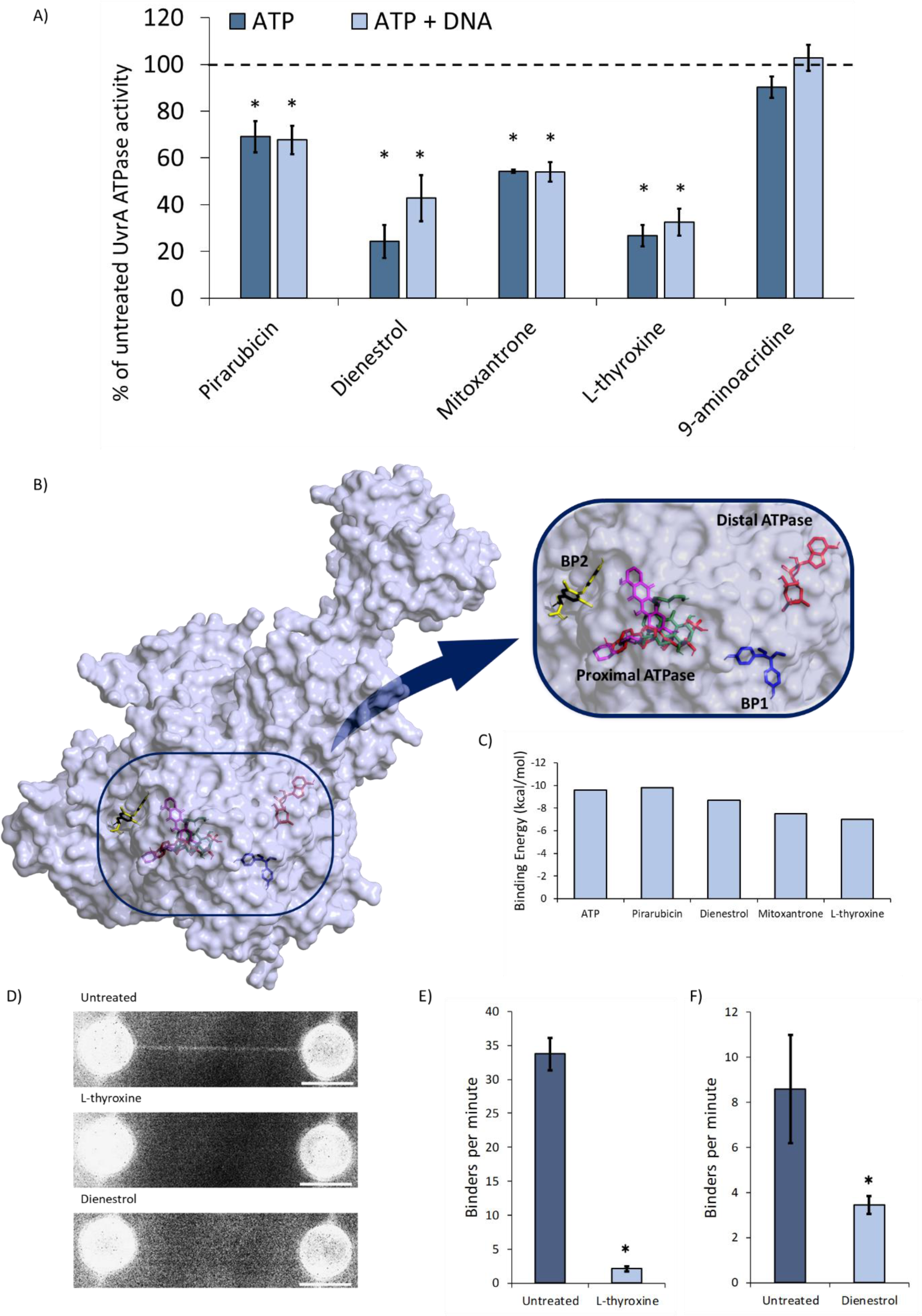
Inhibition of *E. coli* UvrA binding to DNA by selected hits. **A)** NADH-coupled ATPase assay showing the effect of the hits on UvrA’s ATPase activity expressed as percentage of that in the absence of drug (dotted line). The error bars represent the standard error of the mean. Asterisks mark significance: p ≤ 0.05, n = 3 independent replicates. **B)** The Alphafold-calculated structure of the *E. coli* UvrA monomer showing the best docking conformation of the compounds with the greatest effect on UvrA’s ATPase activity. The zoomed in image clearly shows ATP (red), Pirarubicin (magenta), Dienestrol (blue) Mitoxantrone (green), L-thyroxine (yellow) docking. Remarkably L-thyroxine and Dienestrol bind to previously undetected locations on the surface of UvrA. **C)** The minimum binding energy for each compound reveals a range of affinities, although the absolute understanding of these affinities is not clear the values are close or exceed that of ATP (−9.6 kcal/mol). **D)** Using the C-trap binding of UvrA-mNG to a single molecule of DNA could be observed. In the absence of any compounds the average combined fluorescence image from a 10-minute video of DNA shows clear decoration with UvrA (top). In the presence of L-thyroxine (middle) or Dienestrol (bottom) very few molecules bind DNA. **(E & F)** Quantification of DNA binding was provided by the number of binders per minute. This revealed the number of UvrA molecules bound to DNA is significantly reduced in presence L-thyroxine (n=6 strands, p<0.05) or in the presence of Dienestrol (n=6 strands, p<0.05).

To investigate how the most potent ATPase inhibitors could interact with UvrA we performed *in silico* docking using Autodock Vina (27). ATP was used as a control to validate the complete UvrA surface exploration; both of UvrA’s ATP binding sites were successfully located based on comparison with crystal structures (28). The docking precision was underlined by specific interactions being identified with residues K37 (proximal ATP site) and K646 (distal ATP site); these residues have previously been shown to be essential to UvrA’s ATPase activity (23). Each docking predicted a binding energy for ATP of -9.2 kcal/mol and -9.6 kcal/mol, at the proximal and distal sites, respectively. The higher binding affinity of ATP predicted for the C-terminal site confirms the outcomes from a recent study (29), again validating the approach. The strongest affinities are shown as binding energies for each compound in Figure 4C (further data can be found in Table S1). Two compounds showed interaction only with the ATP binding sites; Pirarubicin, which had a stronger affinity than ATP at the proximal site (−9.8 kcal/mol), and Mitoxantrone, which had a preference for the proximal site over the distal, although the binding energy of -7.5 kcal/mol was lower than that of ATP. The docking also revealed two previously unidentified allosteric binding pockets (Figure 4B). Allosteric site ‘BP1’ bound Dienestrol strongly (−8.7 kcal/mol), and the second allosteric site ‘BP2’ close to the proximal ATPase cassette bound L-thyroxine (−7.0 kcal/mol).

To understand if the allosteric binding sites directly affected UvrA binding to DNA we turned to single molecule visualization. Based on our previous data indicating C-terminal fusion of a fluorescent protein to UvrA does not affect function (24), we constructed and expressed C-terminally fused UvrA-mNeonGreen (UvrA-mNG). Both L-thyroxine and Dienestrol bind to allosteric sites and have the strongest reduction in ATPase, and unlike Mitoxantrone and Pirarubicin have not been identified as DNA intercalators (30, 31). In this assay, we suspended a single molecule of DNA between two beads caught in optical traps using the Lumicks C-trap system. Using microfluidics, we established a stream of UvrA alone or UvrA with drug, and the DNA was moved between these streams using the laser tweezers. In the absence of drug, UvrA binds well to the DNA (Figure 4A top), however with either 20 μM L-thyroxine or 20 μM Dienestrol we observed a huge reduction in UvrA binding to the DNA (Figure 4A middle and bottom). Quantification of these interactions by measuring the number of binders per minute over a 10-minute acquisition enabled us to demonstrate a statistically significant reduction in binding.

### Activity against a clinically relevant multi-drug resistant *E. coli* strain

We determined if our successful hits exhibited antibacterial activity against the multidrug-resistant urosepsis-causing *E. coli* clinical isolate, EC958 (32). Since EC958 retains an active TolC pump we only used those compounds effective against WT MG1655. Cisplatin’s MIC (Figure 5A) was identical to that previously reported for WT MG1655 (19), further supporting the use of this approach against multi-drug resistant bacteria. Remarkably, all three drugs showed enhanced activity with cisplatin, for Pirarubicin we observed a 16-fold enhancement of cisplatin MIC and 8-fold for the drug itself. 9-aminoacridine and Mitoxantrone showed an equal improvement for both cisplatin and drugs of 2-fold and 4-fold respectively (figure 5B).

**Figure 5:**
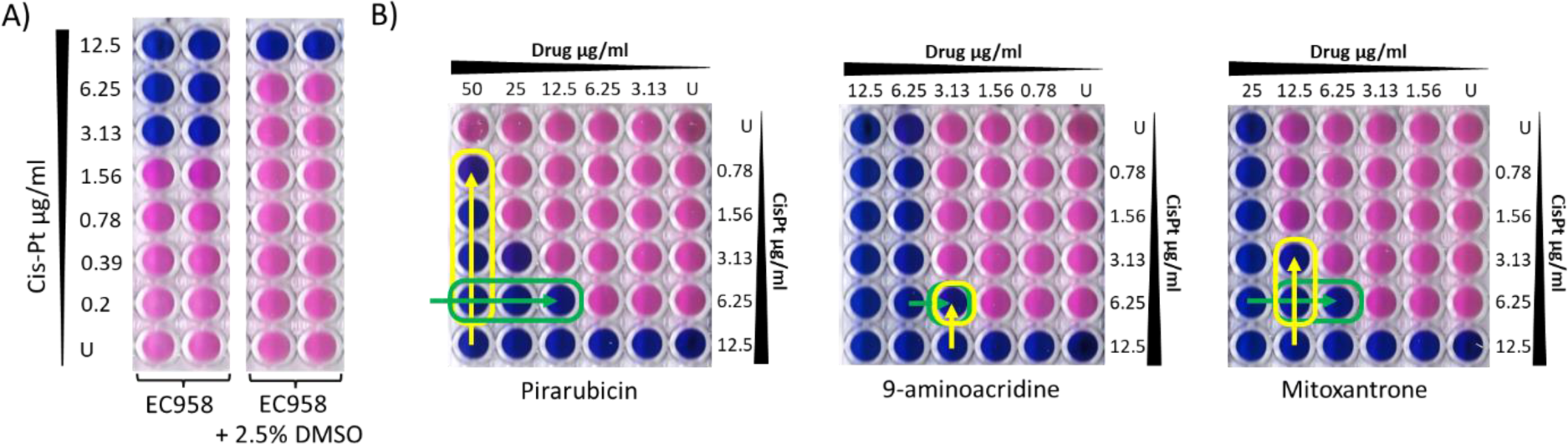
Antibacterial activity of Pirarubicin, 9-aminoacridine and Mitoxantrone against the urosepsis-causing isolate, EC958. As in figure 2 we determined MIC values and performed checkerboard assays. **A)** The cisplatin MIC for EC958 +/- DMSO was determined, and shows DMSO protects EC958 cells from cisplatin. **B)** Checkerboards in EC958 showing the decrease in MIC when combined, the yellow line indicates 2-fold improvement in cisplatin activity per well, the green line indicates 2-fold improvement in drug activity per well. Each plate was replicated three times and these are representative of the repeats. The checkerboards show clear activity and synergy with cisplatin for all of the tested compounds. The panel was limited to these drugs because of their ability to evade the efflux pump TolC.

## Discussion

Adjuvant anti-microbials offer a unique approach to tackle infection in immunocompromised cancer chemotherapy patients. Particularly concerning is the abundance of nosocomial infection as patients with a high potential susceptibility to infection are admitted to hospital. In this study we have developed an approach to the discovery and testing of adjuvant anti-microbials that only possess anti-bacterial activity in the presence of the cancer chemotherapy drug cisplatin. The anti-microbials were sought to target nucleotide excision DNA repair and using a number of evaluation stages, our approach has yielded five final compounds, three of which were shown to possess activity against the *E. coli* clinical isolate, EC958.

All the compounds were validated to target NER both *in vivo* and *in vitro*, providing strong evidence that they function to impair NER in bacteria, allowing cisplatin to kill the cells. To date, there have been a very limited number of studies aimed towards the development of NER inhibitors in bacteria. Of these, a seminal 2011 study screened ∼40k compounds from a general library for effects against mycobacterial NER, using UV as the adjuvant (33). This study identified a single effective compound, however its clinical application is limited because of its poor solubility (20), and problems with using UV as the adjuvant. Therefore, using cisplatin as the DNA damaging agent and screening with an FDA-approved library offers the potential to rapidly progress to the clinic. The compounds thus discovered are currently used in a number of applications ranging from endocrinology to antineoplastic agents.

The latter application is not surprising, since the overlap between antimicrobials and antineoplastics has been well established due to the intention to kill rapidly dividing cells (34, 35). As a consequence, this raises the tantalizing prospect that simply changing the anti-cancer drug treatment regimen might have immediate benefits to patients in terms of reducing infection.

The three lead compounds used against clinical isolates offer the possibility of rapid advancement to the clinic. We also showed these compounds can traverse the bacterial cell membrane and are not efficiently removed by the drug efflux pump, TolC. 9-aminoacridine is used as an externally applied antiseptic, however its use as an antineoplastic agent has recently been proposed due to its action on PI3K (36). Interestingly, 9-aminoacridine has also been used to derivatize cisplatin for improved DNA damaging capabilities (37). It is therefore possible that 9-aminoacridine functions with cisplatin to severely damage the DNA, which overwhelms NER. This would be consistent with the lack of effect on UvrA’s ATPase, however, the clear reduction in incision could equally derive from effects on the other NER proteins. Both Mitoxantrone and Pirarubicin are antineoplastic topoisomerase inhibitors and the mechanism of action for these compounds includes DNA intercalation, although the anthracycline Pirarubicin additionally functions through the generation of reactive oxygen species (38). Although it is easy to imagine the effective drug properties of these compounds mediates through DNA intercalation, we demonstrated that both Mitoxantrone and Pirarubicin directly inhibit UvrA’s ATPase activity in the presence and absence of DNA. The latter point is extremely important, since inhibition is seen without DNA, indicating that intercalation cannot be the sole mechanism of action. Furthermore, we found no inhibition of NER using a Mitoxantrone analogue and members of the Camptothecin family which intercalate DNA (Figure S3).

Using *in silico* docking we were also able to show direct interactions between the compounds and UvrA. Interestingly, two of the compounds, Dienestrol and L-thyroxine had strong affinity for two binding pockets distinct from the ATPase cassette for which they only possessed moderate affinity compared with ATP. These compounds reduced the ATPase activity of UvrA by >70% in the absence of DNA and ∼60% in the presence of DNA; using single molecule imaging it was possible to show this led to severely disrupted DNA binding. These previously unidentified allosteric binding pockets offer potentially new druggable targets on UvrA, and investigations into these new sites are on-going. In this study, we found that L-thyroxine and Dienestrol, two compounds that are not known to intercalate or inhibit bacterial growth, could impair the survival of bacteria when combined with cisplatin and UV radiation. This finding indicates that these compounds may have some synergistic effects with these treatments and opens up the possibility of finding more active analogues based on their chemical scaffold.

The incidence of infection in cancer patients is significantly elevated due to neutropenia and exacerbated by time spent in hospitals leading to nosocomial infection (39, 40). Since the administration of drugs in this study requires the presence of cisplatin, this limits the period over when the drugs will be active. The pharmacokinetics of both drugs will define the therapeutic window, however, the synergy of the combination, as shown in the checkerboard assays, means that lower than MIC drug concentrations are required. This has the effect of lengthening the therapeutic window because as the drugs are excreted, they are still effective at lower concentrations. At present, we are engaged with further understanding the combined pharmacokinetics as a precursor to clinical trial.

In summary, here we have developed a screening strategy to find existing compounds that work in combination with the anticancer therapeutic, cisplatin, which opens up huge potential for the development of new antimicrobials. Screening in combination with other DNA damaging agents will develop NER as a target; potentially offering a much-needed new class of antimicrobial.

## Acknowledgements

We would like to thank the Kad lab for discussions, and Dr James Leech for the initial development of the UV damage repair assay. We also thank Dr Mark Shepherd (University of Kent) for generously providing the EC958 strain.

## Supplementary material

Additional tables and figures are available in the supplementary material, in addition to the materials and methods.

## Notes

### Competing Interest Statement

The authors have declared no competing interest.

## References

1. B. A. Chabner, T. G. Roberts Jr, Timeline: Chemotherapy and the war on cancer. Nat. Rev. Cancer 5, 65–72 (2005).

2. J. Crawford, D. C. Dale, G. H. Lyman, Chemotherapy-induced neutropenia: risks, consequences, and new directions for its management. Cancer 100, 228–237 (2004).

3. K. V. I. Rolston, Infections in Cancer Patients with Solid Tumors: A Review. Infect Dis Ther 6, 69– 83 (2017).

4. T. R. Zembower, Epidemiology of infections in cancer patients. Cancer Treat. Res. 161, 43–89 (2014).

5. A. K. Nanayakkara, et al., Antibiotic resistance in the patient with cancer: Escalating challenges and paths forward. CA Cancer J. Clin. 71, 488–504 (2021).

6. H. C. Neu, The crisis in antibiotic resistance. Science 257, 1064–1073 (1992).

7. F. C. Tenover, Mechanisms of antimicrobial resistance in bacteria. Am. J. Med. 119, S3–10; discussion S62–70 (2006).

8. D. Austin, et al., “Research and Development in the Pharmaceutical Industry” (Congressional Budget Office, 2021) (July 12, 2022).

9. C. Årdal, et al., Antibiotic development - economic, regulatory and societal challenges. Nat. Rev. Microbiol. 18, 267–274 (2020).

10. J. O’Neill, “Tackling Drug-Resistant Infections Globally: Final Report and Recommendations” (London, UK: Review on Antimicrobial Resistance, 2016).

11. B. Rosenberg, L. Van Camp, T. Krigas, Inhibition of cell division in Escherichia coli by electrolysis products from a platinum electrode. Nature 205, 698–699 (1965).

12. S. Hashimoto, H. Anai, K. Hanada, Mechanisms of interstrand DNA crosslink repair and human disorders. Genes Environ. 38, 9 (2016).

13. S. Dasari, P. Bernard Tchounwou, Cisplatin in cancer therapy: Molecular mechanisms of action. European Journal of Pharmacology 740, 364–378 (2014).

14. D. J. Beck, S. Popoff, A. Sancar, W. D. Rupp, Reactions of the UVRABC excision nuclease with DNA damaged by diamminedichloroplatinum(II). Nucleic Acids Res. 13, 7395–7412 (1985).

15. D. Wang, R. Hara, G. Singh, A. Sancar, S. J. Lippard, Nucleotide excision repair from site-specifically platinum-modified nucleosomes. Biochemistry 42, 6747–6753 (2003).

16. C. Petit, A. Sancar, Nucleotide excision repair: from E. coli to man. Biochimie 81, 15–25 (1999).

17. J. J. Truglio, D. L. Croteau, B. Van Houten, C. Kisker, Prokaryotic nucleotide excision repair: the UvrABC system. Chemical Reviews 106, 233–252 (2006).

18. N. M. Kad, B. Van Houten, “Chapter 1 - Dynamics of Lesion Processing by Bacterial Nucleotide Excision Repair Proteins” in Progress in Molecular Biology and Translational Science, P. W. Doetsch, Ed. (Academic Press, 2012), pp. 1–24.

19. A. Gupta, L. Bernacchia, N. M. Kad, Culture media, DMSO and efflux affect the antibacterial activity of cisplatin and oxaliplatin. Lett. Appl. Microbiol. (2022) https://doi.org/10.1111/lam.13767.

20. L. Bernacchia, A. Paris, A. R. Gupta, A. A. Moores, N. M. Kad, Identification of the Target and Mode of Action for the Prokaryotic Nucleotide Excision Repair Inhibitor ATBC. Biosci. Rep. (2022) https://doi.org/10.1042/BSR20220403.

21. A. Sancar, W. D. Rupp, A novel repair enzyme: UVRABC excision nuclease of Escherichia coli cuts a DNA strand on both sides of the damaged region. Cell 33, 249–260 (1983).

22. A. Seck, et al., In vitro reconstitution of an efficient nucleotide excision repair system using mesophilic enzymes from Deinococcus radiodurans. Commun Biol 5, 127 (2022).

23. G. M. Myles, J. E. Hearst, A. Sancar, Site-specific mutagenesis of conserved residues within Walker A and B sequences of Escherichia coli UvrA protein. Biochemistry 30, 3824–3834 (1991).

24. J. T. Barnett, N. M. Kad, Understanding the coupling between DNA damage detection and UvrA’s ATPase using bulk and single molecule kinetics. FASEB J. 33, 763–769 (2019).

25. M. Stracy, et al., Single-molecule imaging of UvrA and UvrB recruitment to DNA lesions in living Escherichia coli. Nat. Commun. 7, 12568 (2016).

26. N. M. Kad, H. Wang, G. G. Kennedy, D. M. Warshaw, B. Van Houten, Collaborative dynamic DNA scanning by nucleotide excision repair proteins investigated by single-molecule imaging of quantum-dot-labeled proteins. Mol. Cell 37, 702–713 (2010).

27. O. Trott, A. J. Olson, AutoDock Vina: improving the speed and accuracy of docking with a new scoring function, efficient optimization, and multithreading. J. Comput. Chem. 31, 455–461 (2010).

28. M. Jaciuk, E. Nowak, K. Skowronek, A. Tañska, M. Nowotny, Structure of UvrA nucleotide excision repair protein in complex with modified DNA. Nat. Struct. Mol. Biol. 18, 191–198 (2011).

29. B. C. Case, S. Hartley, M. Osuga, D. Jeruzalmi, M. M. Hingorani, The ATPase mechanism of UvrA2 reveals the distinct roles of proximal and distal ATPase sites in nucleotide excision repair. Nucleic Acids Res. 47, 4136–4152 (2019).

30. P. J. Smith, S. A. Morgan, M. E. Fox, J. V. Watson, Mitoxantrone-DNA binding and the induction of topoisomerase II associated DNA damage in multi-drug resistant small cell lung cancer cells. Biochem. Pharmacol. 40, 2069–2078 (1990).

31. H.-Y. Zou, et al., Studying the interaction of pirarubicin with DNA and determining pirarubicin in human urine samples: combining excitation-emission fluorescence matrices with second-order calibration methods. J. Fluoresc. 19, 955–966 (2009).

32. C. A. Ribeiro, et al., Nitric oxide (NO) elicits aminoglycoside tolerance in Escherichia coli but antibiotic resistance gene carriage and NO sensitivity have not co-evolved. Arch. Microbiol. 203, 2541–2550 (2021).

33. N. Mazloum, et al., Identification of a chemical that inhibits the mycobacterial UvrABC complex in nucleotide excision repair. Biochemistry 50, 1329–1335 (2011).

34. V. W. C. Soo, et al., Repurposing of Anticancer Drugs for the Treatment of Bacterial Infections. Curr. Top. Med. Chem. 17, 1157–1176 (2017).

35. C. Herold, et al., Ciprofloxacin induces apoptosis and inhibits proliferation of human colorectal carcinoma cells. Br. J. Cancer 86, 443–448 (2002).

36. C. Guo, et al., 9-Aminoacridine-based anticancer drugs target the PI3K/AKT/mTOR, NF-kappaB and p53 pathways. Oncogene 28, 1151–1161 (2009).

37. H. W. Kava, W. Y. Leung, A. M. Galea, V. Murray, The DNA binding properties of 9-aminoacridine carboxamide Pt complexes. Bioorg. Med. Chem. 40, 116191 (2021).

38. G. N. Hortobágyi, Anthracyclines in the treatment of cancer. An overview. Drugs 54 Suppl 4, 1–7 (1997).

39. M. Kamboj, K. A. Sepkowitz, Nosocomial infections in patients with cancer. Lancet Oncol. 10, 589–597 (2009).

40. A.-M. Jiang, et al., Nosocomial infections due to multidrug-resistant bacteria in cancer patients: a six-year retrospective study of an oncology Center in Western China. BMC Infect. Dis. 20, 452 (2020).

