## Supplementary information for "Combining cancer chemotherapeutics with bacterial DNA repair inhibitors to develop novel antimicrobials"

**Table S1:** Calculated binding energies of compounds to UvrA’s ATP binding pockets and allosteric sites

| Compound | Binding energy (kcal/mol)* | | |
| --- | --- | --- | --- |
| ATP | -9.6 Distal | -9.2 Proximal |  |
| Pirarubicin | -9.8 Proximal | -9.6 Distal |  |
| Dienestrol | -8.7 Allosteric BP1 | -7.4 Distal |  |
| Mitoxantrone | -7.5 Proximal | -7.4 Distal |  |
| L-Thyroxine | -7.0 Allosteric BP2 | -6.9 Distal | -6.9 Proximal |

**Distal refers to C-terminal ATP binding pocket, proximal refers to N-terminal binding pocket, Allosteric BP (1,2) refer to newly determine allosteric sites located on the surface of UvrA (see Figure 4B).*

**UV damage repair assay controls**

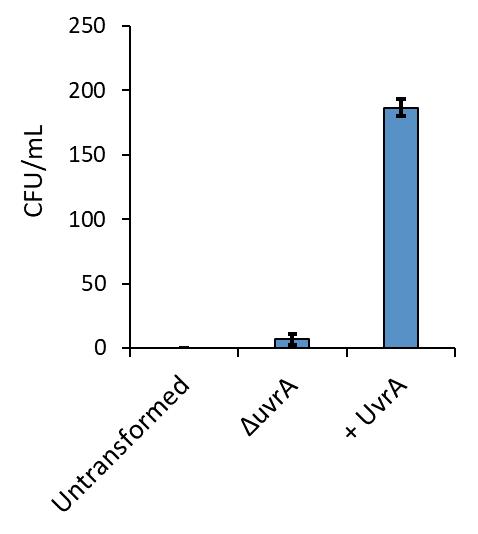

**Figure S1: UV damage repair assay controls.** The colony forming units (CFUs) of MG1655 spread on ampicillin plates in the absence of any introduced plasmid (untransformed), or when UvrA is knocked-out (*ΔuvrA*) show no growth. In the presence of functioning NER (+UvrA) the damage is efficiently repaired, conferring resistance, leading to substantial growth. Error bars represent the standard error of the mean (n≥3).

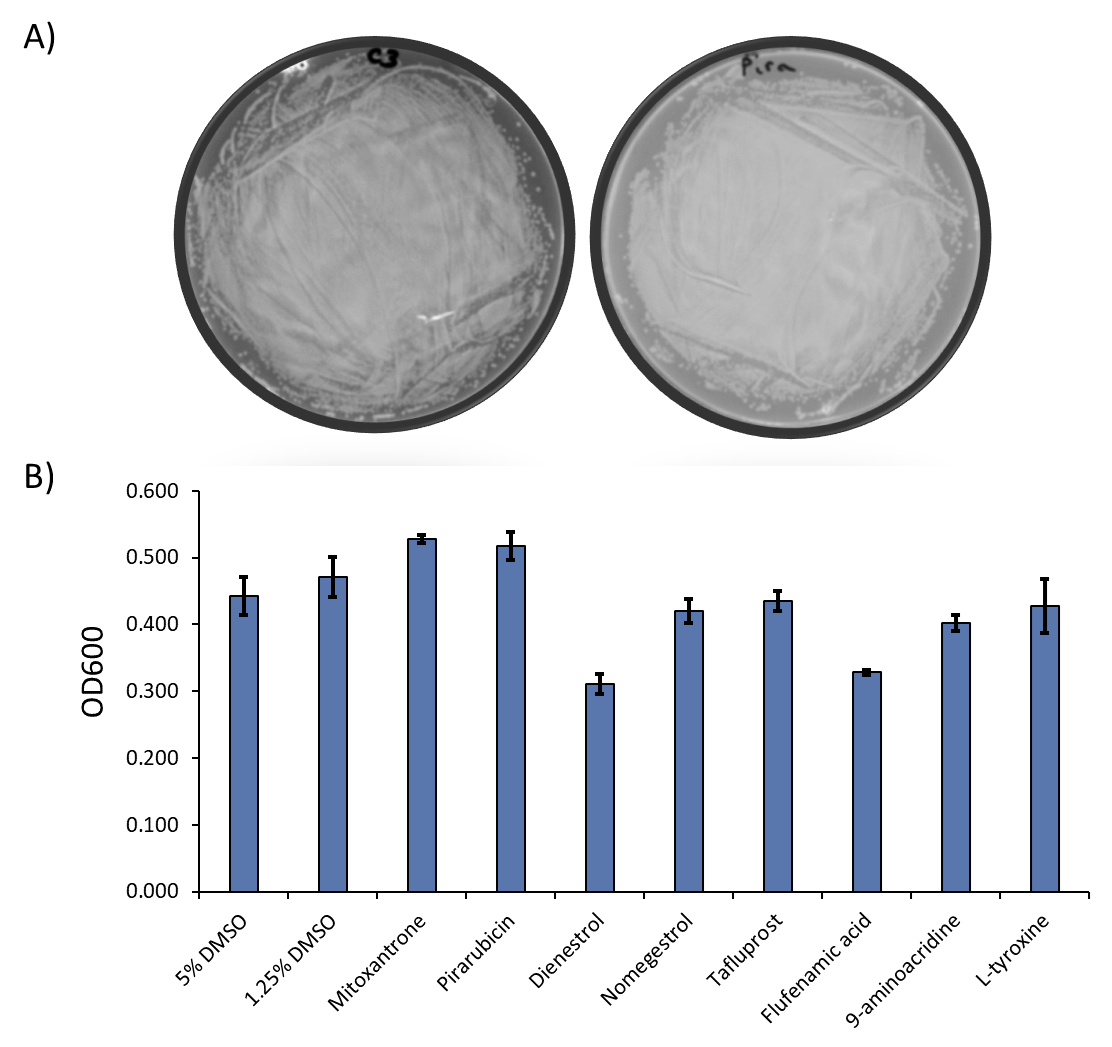

**Figure S2: Compounds alone do not impair growth in the UV growth repair assay.** A) Example plate showing MG1655 spread on LB agar in the absence of added drug compounds (left) and in the presence of the same concentration of Pirarubicin used in figure 3D (right). No effect on growth is seen. The plates are representative of three different replicates for each of the compounds. B) OD_600_ of the same cultures shows a minimal decrease in cells density when incubated with the compounds. The largest effect measured of approximately 35% and 25% for Dienestrol and Flufenamic acid respectively is not large enough to explain the complete lack of growth in the UV damage repair assay.

**Intercalation controls**

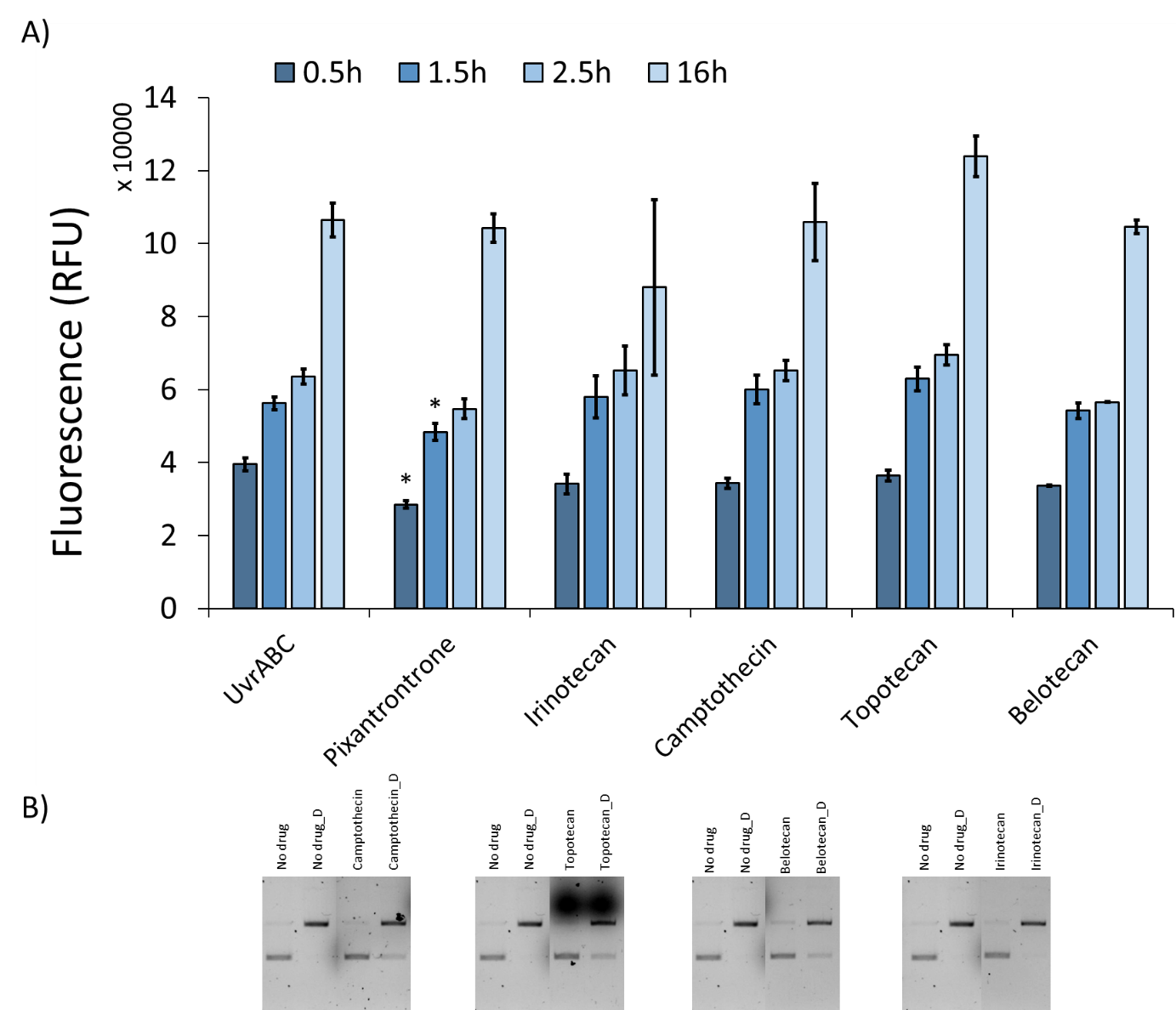

**Figure S3: Known intercalators cannot inhibit the incision mediated by NER. A) Known intercalators tested for their anti NER activity using the fluorescence incision assay**. Pixantrone is a DNA intercalator (1) analogue to Mitoxantrone which resulted able to drastically inhibit incision at 16h. Pixantrone however is unable to considerably delay incision when tested in the fluorescence-base assay. The error bars represent the standard error of the mean (n≥2). The result were considered significant when compared to the untreated control (UvrABC) when p ≥ 0.05 (*). Irinotecan, Camptothecin, Topotecan and Belotecan are also known DNA intercalators (2, 3) unable to significantly alter the incision when tested. B) Gel-based incision assay performed as further confirmation on the some of the compounds available. The results confirm what shown with the fluorescence based incision assay, the intercalating action alone is not the primary responsible for UvrABC inhibition. The pictures are representative of two independent replicates.

**Materials and methods**

**Bacterial strains, media, and culture conditions**

The strains used in this study include *E. coli* MG1655, MG1655 Δ*uvrA,* MG1655 Δ*tolC,* MG1655 Δ*uvrA* Δ*tolC*, BL21 Δ*uvrA* Δ*uvrB* and EC958 (4) . The knockout strains were previously generated (5) using P1 transduction of the respective gene deletions from the Keio collection (6). Luria Bertani broth and agar was used for primary culture and maintenance of bacterial strains. Strains were sub-cultured in Mueller Hinton Broth II (MHBII, Sigma, Dorset, UK) or MOPS (Melford, Berkshire, UK) minimal medium (pH 7.4) (7)  supplemented with 0.2% glucose, 1.32 mM K_2_HPO_4_ and 0.1 μg/ml thiamine.  All strains were grown at 37^o^C with vigorous shaking.

**Antimicrobials and chemicals**

The library of 2731 FDA-approved drugs were purchased from MedChemExpress, cisplatin and 9-aminoacridine were purchased from Sigma, Dorset, UK. All other compounds used for testing antimicrobial activity were purchased from MedChemExpress.

**Cloning, expression and purification of *E. coli* UvrA, UvrB and UvrC**

Unlabelled UvrB and UvrC (amplified from *E coli* MG1655, NC_0009313.3) were cloned into the IPTG inducible vector pJB using primer pairs - UvrB.gibson_F (5’-ATGAGATCCTCTCATAGTTAATTTC-3’) and UvrB.gibson_R (5’-GGCGCGCCTTCAGGTAGC-3’) for UvrB and UvrC.gibson_F (5’-ATGAGATCCTCTCATAGTTAATTTC-3’) and UvrC.gibson_R (5’-GGCGCGCCTTCAGGTAGC-3’) for UvrC. Proteins were engineered with a flexible C-terminal linker, followed by an AviTag, TEV protease site and 6X His-tag for purification using a Ni-NTA column. The unlabelled and mNeonGreen-tagged UvrA used in this study were expressed, purified and stored as described previously (5), with the exception that the salt concentration was increased to 100 mM KCl and 400 mM KCl for the storage of UvrB and UvrC, respectively.

**High-throughput screening**

The FDA-approved library of 2731 compounds were screened against MG1655 and MG1655 Δ*tolC* in the presence of 4 μg/mL cisplatin in MOPS minimal medium (7) as described in Figure 1A. The compounds were screened at a final concentration of 20 μM and resazurin was used to determine growth inhibition (Figure 1B). After initial screening, compounds exhibiting growth inhibition in the presence of cisplatin were screened in the presence of UV at 75 J/m^2^ at 254 nm using a UV cross-linker (UVP/Analytik Jena UV Crosslinker CX-2000). Compounds which retained antibacterial activity in the presence of 75 J/m^2^ UV were moved forwards and those that did not show synergy in the presence of UV irradiation were kept aside for future characterisation. The shortlisted compounds were subsequently characterised as described in Figure 1A.

**Antibacterial susceptibility testing**

The minimum inhibitory concentration of antimicrobial compounds used in this study was determined following CLSI guidelines (8) against MG1655, MG1655 Δ*uvrA*, MG1655 Δ*tolC*, and MG1655 Δ*uvrA* Δ*tolC,* in MOPS minimal medium (7). Resazurin was used to determine the MIC as described previously (5).

**Checkerboard assay**

The checkerboard assay was used to assess activity of test compounds in the presence of cisplatin. The broth microdilution-based checkerboard assay was used with a few modifications (9, 10). A 2-fold series dilution of drugs in DMSO were plated across the plates ([DMSO] =2.5%). Subsequently a 2-fold dilution of cisplatin in 0.9 % v/v saline was plated (2.5 µl) in the microplate carefully without mixing the two solutions. Finally, 95 μL of a bacterial suspension in MOPS minimal medium prepared following CLSI guidelines was added to the wells (7, 8) to ensure activity of cisplatin (11).

**NADH-linked ATPase Assay**

This assay was modified from the previously described protocol (5) to enable larger numbers of compounds to be assessed in a 96-well plate format. A 50 nM UvrA master mix (50 mM Tris (pH 7.5), 50 mM KCl, 10 mM MgCl_2_, 0.5 mM phosphoenolpyruvate, 1 mM DTT, 210 μM NADH, 2% v/v pyruvate kinase (600–1000 U/ml) and lactate dehydrogenase (900–1400 U/ml, premixed stock from Merck)) was prepared for each independent replicate. Each reaction well consisted of a 200 µL of the above master mix and 20 µM of drug. In addition, an untreated control containing 200 µL 50 nM UvrA master mix and 2.5% DMSO were performed per plate, this yielded the basal activity of UvrA. The components were incubated at room temperature for ~5 minutes and then the reaction was started with the addition of 1 mM ATP. After ~6 minutes 50 ng pUC18 DNA was added to the reaction mix. All results were reported as a percentage of this basal activity. The experiments were performed in triplicate and the error bars represent the standard error of the mean.

**Agarose gel-based DNA incision assay**

Each incision reaction was carried out in 20 μL final volume of ABC Buffer (50 mM Tris (pH 7.5), 50 mM KCl, 10 mM MgCl_2_, 0.0001% sodium azide) containing 5 mM DTT, 1 mM ATP, 100 nM UvrA, 200 nM UvrB, 100 nM UvrC, 28 nM of pUC18 DNA irradiated at 200 J/m^2^ at 254 nm in the UV crosslinker (or undamaged DNA), and 20 μM of compound when indicated. The mixtures were incubated at 37^o^C for 30 mins and the reaction was stopped by heat inactivation at 65^o^C for 10 mins. DNA was separated and visualised on a 0.8% agarose gel.

**Fluorescence-based incision assay**

Two complementary oligonucleotides were designed to incorporate a fluorescein modified thymine as a substrate for NER (12). /BHQ/GT AAC TAA GCT TGA CGA TGG AGC CGT AAC AGT ACG TAG TCT G and CAG ACT ACG TAC TGT TAC GGC TCC ATC /**F**T/TC AAG CTT AGT TAC /Cy5/ (Integrated DNA Technologies, IDT, UK). At the 3’ terminus of the damage (fluorescein) containing strand we placed a Cy5 fluorophore, and the 5’ terminus of the complementary strand was labelled with an Iowa black hole quencher (BHQ) to quench Cy5 fluorescence when annealed. UvrC incision of the fluorescein containing oligonucleotide (14 bases on 3` and 27 bases on 5`) would produce a 10-base long oligonucleotide with a melting temperature of approximately 16^o^C, leading to an increase in fluorescence. The total volume per reaction in each well of a 384-well fluorescence plate was 25 µl. Each reaction mix contained 5 mM DTT, 200 nM UvrA, 400 nM UvrB, 200 nM UvrC, and 1 mM ATP in ABC buffer and was prepared on ice. 100 nM of the DNA substrate, pre-annealed by heating to 95^o^C for 5 minutes and then slow cooling to room temperature was added to initiate the reaction. Cy5 fluorescence readings were taken from the bottom after 0.5/1.5/2.5/16 h at 37^o^C. The outer wells were filled with water to generate a humid environment in the plate to minimize evaporation. The experiments were repeated at least twice with 2 technical replicates each, the error was reported as the standard error of the mean (n≥4).

**Repair assay for a UV damaged plasmid**

Chemically competent cells of MG1655 Δ*tolC* and MG1655 Δ*uvrA* Δ*tolC* were prepared according to the method previously described (13) and stored at -80^o^C. Prior to transformation 50 µl of cells were thawed on ice for 20 minutes before 100 ng of pUC18 DNA, irradiated at 200 J/m^2^ at 254 nm, was added and incubated on ice for 30 minutes. Subsequently, cells were incubated at 42^o^C for 30 seconds and then immediately placed on ice for two minutes. Cells were then transferred to 350 µl LB broth containing 2x MIC of selected antimicrobial agents and incubated for an hour at 37^o^C with aeration. Cells were pelleted and resuspended in pre-warmed 350 µl LB. Aliquots where then plated in LB agar containing ampicillin (100 µg/mL) and incubated at 37^o^C overnight.

To ensure that growth inhibition in this repair assay was due to activity against NER instead of the compounds directly killing the bacteria, an additional control was performed. Cells were grown to 0.5 (OD_600_) from an overnight culture and after pelleting were resuspended in LB to a volume commensurate with the same cell concentration present in the UV damage repair assay. These cells were incubated for one hour in presence or absence of drug and at the end of the incubation period the bacteria were pelleted and resuspended in fresh LB broth. 5 µl were then streaked onto an LB agar plate and incubated at 37^o^C overnight. In parallel, 100 µl of the cell suspension was transferred into a microtiter plate for OD_600_ measurement.

***In silico* drug docking**

To study possible binding of the selected compounds to UvrA computational docking of UvrA was performed following the adapted protocol described previously (5). Since no *E. coli* UvrA structure is available *E. coli* UvrA’s protein structure was retrieved from the AlphaFold database (14, 15), and then converted into pdbqt using Autodock tools 1.5.7 (16). The 3D structures of the compounds were downloaded from PubChem or Zinc databases (17, 18), energy minimized, and converted into the pdbqt format using OpenBabel (19). Autodock Vina (20) was used to explore the possible binding of the most interesting compounds to UvrA. The search space was maximised to include the entirety of the protein and the exhaustiveness increased 1000 times (to 8000 from the standard setting of 8) to minimize the effect of a large search space and thus obtaining more accurate docking. As a control, ATP was docked and found to bind the distal and proximal ATPase cassettes validating the ability of the algorithm to find reliable results. Both the docking models and the binding energy were analysed to find possible binding sites on the protein and hypothesize a mechanism of action.

**Single molecule microscopy**

To directly visualize protein binding to DNA we used optical tweezers coupled with fluorescence imaging (C-trap, Lumicks, NL). This system uses microfluidics to allow the capture of a single end-biotinylated Lambda DNA molecule between two streptavidin-coated silica beads. To visualise UvrA-mNeonGreen binding to DNA we transformed a plasmid containing the C-terminally tagged UvrA-mNeonGreen into BL21 Δ*uvrA* Δ*uvrB* (5) and grew the cells to mid-log phase (0.4-0.6 OD600) before induction of expression with 0.5 mM IPTG at 37 °C for 3 hours. Cells were spun at 20000 rpm for 30 minutes at 4^o^C and resuspended in buffer (50 mM NaH_2_PO_4_, 500 mM NaCl, 15 mM imidazole pH 8). 100 µg/ml lysozyme was used with sonication to ensure complete cell lysis in the presence of protease inhibitor cocktail (no EDTA) (ThermoFisher) and 1mM PMSF. The cell debris were spun from solution at 20000 rpm for 30 mins at 4^o^C, and the concentration of UvrA-mNeonGreen in the supernatant was determined by absorption at 506 nm (extinction coefficient = 116000 M^-1^cm^-1^). Prior to imaging, the lysate was diluted in ABC buffer supplemented with 5 mM DTT and 1 mM ATP to a final UvrA-mNeonGreen concentration of 5nM. Finally, the solution was clarified using a 0.22 μm syringe filter before being applied to the system.

Using the microfluidics capacity of the C-trap, we recorded UvrA-mNeonGreen binding to DNA in a channel with no compound present (untreated), followed by measuring binding to the same strand of DNA in the compound-containing channel (treated), and vice versa (n=6 strands total). Each video was recorded for 10 minutes with a 200 ms exposure at a framerate of 2 Hz using exposure synchronisation. Videos were analysed using the TrackMate plugin of Fiji (ImageJ), to objectively count the number of binders.

**References**

1.	D. Mukherji, R. Pettengell, Pixantrone for the treatment of aggressive non-Hodgkin lymphoma. *Expert Opin. Pharmacother.* **11**, 1915–1923 (2010).

2.	Y. Temerk, M. Ibrahim, H. Ibrahim, W. Schuhmann, Comparative studies on the interaction of anticancer drug irinotecan with dsDNA and ssDNA. *RSC Adv.* **8**, 25387–25395 (2018).

3.	B. L. Staker, *et al.*, The mechanism of topoisomerase I poisoning by a camptothecin analog. *Proc. Natl. Acad. Sci. U. S. A.* **99**, 15387–15392 (2002).

4.	C. A. Ribeiro, *et al.*, Nitric oxide (NO) elicits aminoglycoside tolerance in Escherichia coli but antibiotic resistance gene carriage and NO sensitivity have not co-evolved. *Arch. Microbiol.* **203**, 2541–2550 (2021).

5.	L. Bernacchia, A. Paris, A. R. Gupta, A. A. Moores, N. M. Kad, Identification of the Target and Mode of Action for the Prokaryotic Nucleotide Excision Repair Inhibitor ATBC. *Biosci. Rep.* (2022) https:/doi.org/10.1042/BSR20220403.

6.	T. Baba, *et al.*, Construction of Escherichia coli K-12 in-frame, single-gene knockout mutants: The Keio collection. *Mol. Syst. Biol.* **2**, 2006.0008 (2006).

7.	F. C. Neidhardt, P. L. Bloch, D. F. Smith, Culture medium for enterobacteria. *J. Bacteriol.* **119**, 736–747 (1974).

8.	F. R. Cockerill, *et al.*, Methods for Dilution Antimicrobial Susceptibility tests for bacteria that grow Aerobically: Approved Standard-ninth Edition. *CLSI* **32** (2012).

9.	M. Balouiri, M. Sadiki, S. K. Ibnsouda, Methods for in vitro evaluating antimicrobial activity: A review. *J Pharm Anal* **6**, 71–79 (2016).

10.	M. H. Hsieh, C. M. Yu, V. L. Yu, J. W. Chow, Synergy assessed by checkerboard. A critical analysis. *Diagn. Microbiol. Infect. Dis.* **16**, 343–349 (1993).

11.	A. Gupta, L. Bernacchia, N. M. Kad, Culture media, DMSO and efflux affect the antibacterial activity of cisplatin and oxaliplatin. *Lett. Appl. Microbiol.* (2022) https:/doi.org/10.1111/lam.13767.

12.	M. J. DellaVecchia, *et al.*, Analyzing the Handoff of DNA from UvrA to UvrB Utilizing DNA-Protein Photoaffinity Labeling*♦. *J. Biol. Chem.* **279**, 45245–45256 (2004).

13.	R. Green, E. J. Rogers, Transformation of chemically competent E. coli. *Methods Enzymol.* **529**, 329–336 (2013).

14.	J. Jumper, *et al.*, Highly accurate protein structure prediction with AlphaFold. *Nature* **596**, 583–589 (2021).

15.	M. Varadi, *et al.*, AlphaFold Protein Structure Database: massively expanding the structural coverage of protein-sequence space with high-accuracy models. *Nucleic Acids Res.* **50**, D439–D444 (2022).

16.	S. Forli, *et al.*, Computational protein-ligand docking and virtual drug screening with the AutoDock suite. *Nat. Protoc.* **11**, 905–919 (2016).

17.	J. J. Irwin, *et al.*, ZINC20-A Free Ultralarge-Scale Chemical Database for Ligand Discovery. *J. Chem. Inf. Model.* **60**, 6065–6073 (2020).

18.	S. Kim, *et al.*, PubChem in 2021: new data content and improved web interfaces. *Nucleic Acids Res.* **49**, D1388–D1395 (2021).

19.	N. M. O’Boyle, *et al.*, Open Babel: An open chemical toolbox. *J. Cheminform.* **3**, 33 (2011).

20.	O. Trott, A. J. Olson, AutoDock Vina: improving the speed and accuracy of docking with a new scoring function, efficient optimization, and multithreading. *J. Comput. Chem.* **31**, 455–461 (2010).
